# Gene expression divergence following gene and genome duplications in spatially resolved plant transcriptomes

**DOI:** 10.1101/2025.05.27.656262

**Authors:** Fabricio Almeida-Silva, Yves Van de Peer

**Affiliations:** Department of Plant Biotechnology and Bioinformatics, Ghent University, 9052 Ghent, Belgium; VIB Center for Plant Systems Biology, VIB, 9052 Ghent, Belgium; Centre for Microbial Ecology and Genomics, Department of Biochemistry, Genetics and Microbiology, University of Pretoria, Pretoria 0028, South Africa; College of Horticulture, Academy for Advanced Interdisciplinary Studies, Nanjing Agricultural University, Nanjing, China

**Keywords:** polyploidy, spatial transcriptomics, transcriptional regulation, regulatory genomics

## Abstract

Gene and genome duplications are key drivers of plant genome evolution, expanding genetic repertoires and facilitating functional innovation. However, genes originating from different duplication mechanisms undergo different outcomes, especially at the expression level. Previous studies have demonstrated that segmental or whole-genome duplications generate duplicates with similar and somewhat redundant expression profiles across multiple tissues, while other duplicates display increased divergence, ultimately leading to functional innovations. However, little is known about how duplicates diverge in expression across cell types in a single tissue. Here, we used high-resolution spatial transcriptomic data from five species (*Arabidopsis thaliana, Glycine max, Phalaenopsis aphrodite, Zea mays,* and *Hordeum vulgare*) to investigate the evolution of gene expression following gene duplications. We found that genes originating from segmental or whole-genome duplications display increased expression levels, expression breadths, spatial variability, and number of coexpression partners. Duplication mechanisms that preserve cis-regulatory landscapes typically generate paralogs with more preserved expression profiles, but such differences by duplication mode disappear over time. Expression divergence also depends on gene functions, with dosage-sensitive gene families displaying highly preserved expression profiles, while families involved in more specialized processes (e.g., flowering and phytohormone biosynthesis) display increased divergence. Paralogs originating from large-scale (including whole-genome) duplications display redundant and/or overlapping expression profiles, indicating functional redundancy and/or subfunctionalization, while small-scale duplicates diverge asymmetrically, indicating neofunctionalization. Collectively, our findings provide new insights into the tempo and mode of gene expression evolution, helping understand how gene and genome duplications shape cell identities.

## Introduction

Gene duplications have long been recognized as a major source of novel genetic material upon which evolution can act (Ohno 1970). Individual genes can be duplicated locally (though tandem and proximal duplications), and/or through transposon-mediated duplications (Panchy et al. 2016; Almeida-Silva and Van de Peer 2025). In plants, single-gene duplications have fueled the evolution of key traits, including secondary metabolism and stress response (Baumgarten et al. 2003; Rizzon et al. 2006; Chae et al. 2014; Almeida-Silva and Van de Peer 2023; Ji et al. 2024). Whole-genome duplications (WGDs) are an extreme duplication mechanism whereby all chromosome sets double in copy number, and they have occurred multiple times in all eukaryotic kingdoms, but are particularly pervasive in plants (Van De Peer et al. 2017; Ren et al. 2018; One Thousand Plant Transcriptomes Initiative 2019; Almeida-Silva and Van de Peer 2025). Despite the detrimental effects of doubling the entire genome, such as reduced fertility and genomic instability (Comai 2005; Woodhouse et al. 2010), evidence for the evolutionary and ecological potential of WGD has accumulated steeply, with WGDs being associated with increased diversity (Tank et al. 2015; Landis et al. 2018; Ren et al. 2018), origin of novel traits (Soltis and Soltis 2016), increased robustness in gene regulatory networks (Almeida-Silva and Van de Peer 2023), and survival during environmental catastrophes (Chen et al. 2024).

Duplications of genes and entire genomes immediately result in redundancy, allowing for the accumulation of mutations in coding and *cis*-regulatory sequences through genetic drift. While redundancy can be selected for in special cases (e.g., to increase mutational robustness), duplicated genes are thought to undergo one of three major outcomes in the long term: gene loss (pseudogenization), partitioning of ancestral functions (subfunctionalization), or gain of novel functions (neofunctionalization) (Panchy et al. 2016; Birchler and Yang 2022; Iohannes and Jackson 2023). Importantly, the fate of a duplicated gene largely depends on its duplication mechanism and functions. For instance, dosage-sensitive gene families (e.g. regulatory proteins and multiprotein complexes) are typically lost after single-gene duplications, but not after whole-genome duplications, what is likely due to selection to preserve stoichiometric balance in a cell (Blanc and Wolfe 2004; Maere et al. 2005; Veitia et al. 2008; Freeling 2009; Birchler and Veitia 2010). Likewise, retained defense-related genes often undergo neofunctionalization after tandem and/or proximal duplications (Birchler and Yang 2022; Ugalde and Straube 2023).

Over the past decades, several studies have explored large compendia of expression data to investigate how paralogous genes diverge in expression following gene and genome duplications. For instance, WGD-derived duplicates in *Arabidopsis thaliana* display highly redundant and overlapping expression profiles, while duplicates created by small-scale duplications display asymmetric divergence (Casneuf et al. 2006). Also in *A. thaliana*, WGD-derived genes display greater expression levels and are expressed in more tissues (Ganko et al. 2007). In rice (*Oryza sativa*), segmental and tandem duplicates display higher expression correlations compared with duplicates originating from other modes (Li et al. 2009). In soybean (*Glycine max*), WGD-derived paralogs display greater levels of expression preservation compared to other duplication modes, with particularly larger differences for pairs originating at the *Glycine*-specific WGD at around 13 million years ago (Almeida-Silva et al. 2020a).

Although analyses of large-scale, multi-tissue expression compendia have provided important insights into expression-level divergence following different modes of duplications, they are limited in resolution. As bulk RNA-seq samples aggregate gene expression across multiple (often heterogeneous) cell populations, cell type- and cell state-specific changes in expression following duplications are not captured. While single-cell RNA sequencing offers high-resolution expression profiling, it also comes with limitations, such as loss of original spatial information (important for accurate cell type annotation), and biased proportions for cell types that are rare and/or difficult to protoplast (Kiselev et al. 2019; Xu et al. 2021; Cox Jr et al. 2022; Yin et al. 2023). Spatial transcriptomics addresses such issues, allowing for *in situ* quantification of transcript abundances. Investigating how paralogous genes diverge in expression within tissue compartments using spatially resolved transcriptomes could help understand how gene and genome duplications shape cell identities.

Here, we analyzed spatially resolved plant transcriptomes from five species to investigate how duplicated genes diverge in expression across tissue compartments. We found that genes originating from different duplication modes display differences in expression levels, expression breadth, and spatial expression variability. Expression divergence between paralogous gene pairs increases exponentially with time, and varies in an age- and duplication-mode specific manner. Expression divergence varies across gene families, with dosage-dependent gene families displaying greater preservation. Segmental duplicates are highly connected in coexpression networks and typically display expression-level redundancy or symmetric divergence (suggestive of subfunctionalization), while small-scale duplicates (tandem, proximal, and transposed) typically diverge asymmetrically (suggestive of neofunctionalization). Collectively, our findings provide important insights into the impact of gene and genome duplications on plant cell identities.

## Results

### Data set overview and sample quality statistics

We manually curated the literature to find plant spatial transcriptomics data sets for which count matrices and spot metadata were publicly available (see Materials and Methods for details). We obtained five independent data sets, which represent Arabidopsis cauline leaves (Xia et al. 2022), soybean (*Glycine max)* nodules infected with rhizobia (Liu et al. 2023), developing maize (*Zea mays*) ears (Wang et al. 2024), *Phalaenopsis* Big Chili inflorescences (buds and shoot apical meristem) at successive developmental stages (hereafter referred to as *Phalaenopsis aphrodite*, whose genome was used as a reference in the original publication) (Liu et al. 2022), and germinating barley *(Hordeum vulgare*) seeds (Peirats-Llobet et al. 2023) (Table 1, Supplementary Fig. S1a).

**Table 1.**
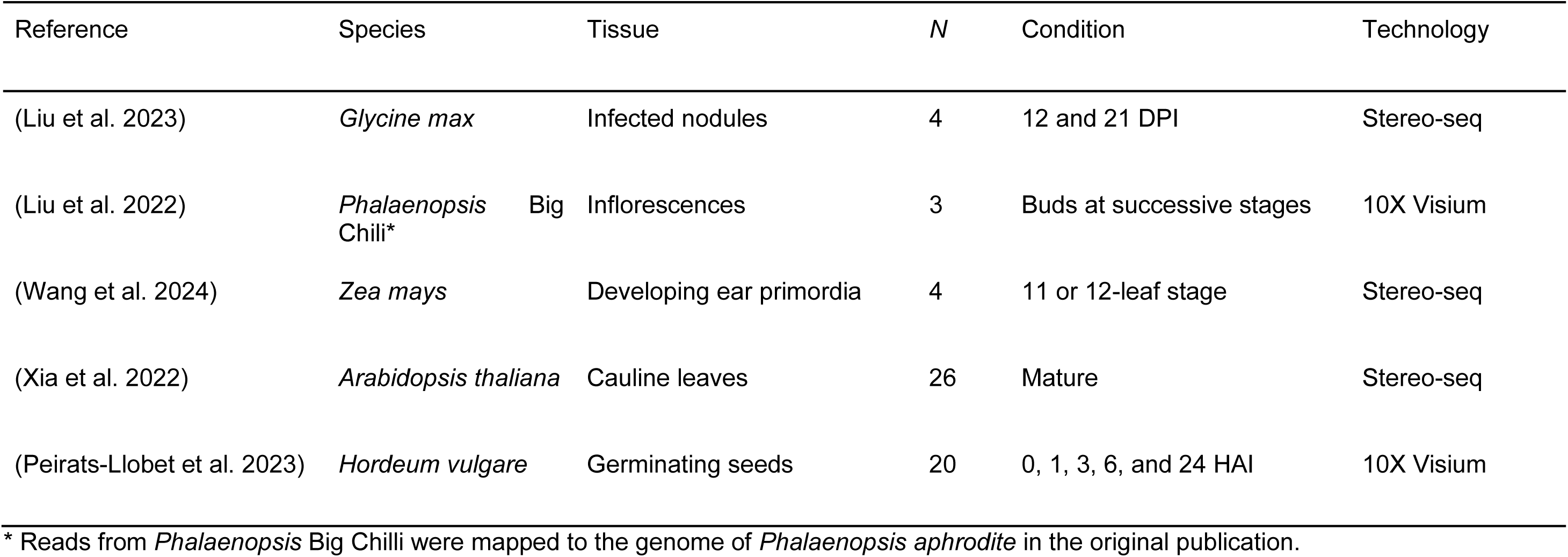
Spatial transcriptomic data sets included in this study. *N*, number of tissue slides. DPI, days post-infection. HAI, hours after imbibition.

We performed minimal preprocessing to remove genes with zero counts across all spots, spots with zero counts, and genes with less than 1 count in ≥99.9% of the spots. After preprocessing, all samples had >10,000 genes, and the median number of spots per sample in each species ranged from 1545 in *H. vulgare* to 8062 in *Zea mays* (Supplementary Fig. S1b). The number of spots per sample in *A. thaliana* was exceptionally smaller than that for other species (median = 631) because of the considerably smaller tissue area in transversal leaf sections (Supplementary Fig. S1a). The median number of distinct spatial domains per sample ranged from 6 in *A. thaliana* and *G. max* to 14 in *H. vulgare*. Collectively, summary gene- and spot-level statistics indicate that samples display high quality, as all samples have had their transcriptomes profiled in a high-throughput manner (over 10,000 genes) across hundreds to thousands of spots.

### Gene duplication mode impacts expression levels, expression breadths, and spatial variability

To investigate whether genes originating from different duplication modes display differences in global expression profiles, we first classified genes as originating from segmental (SD), tandem (TD), proximal (PD), retrotransposed (rTRD), DNA transposed (dTRD), or dispersed duplications (DD) (see Materials and Methods for details). Importantly, SD gene sets include both genes created by WGD and by duplications of large genomic regions (e.g., entire chromosomes, entire chromosome arms, etc), so we chose to broadly use ‘SD’. Then, we tested for differences in mean expression levels, expression breadths (i.e., number of distinct cell types in which genes are expressed), and spatial variability (i.e., variability in expression across spatial domains).

We observed that duplication modes that preserve *cis*-regulatory landscapes (i.e., SD and TD) often generate genes with greater expression levels and expression breadth, especially compared to DD genes (Kruskal-Wallis test, P <0.05; Fig. 1a and 1b). However, dTRD genes in *H. vulgare* and *P. aphrodite* are exceptions, with expression levels and breadths comparable to SD and TD genes. In all species, SD genes are overrepresented in spatially variable gene (SVG) sets, indicating that genes originating from this duplication mode are more likely to display variable expression profiles across tissue compartments (Fisher’s exact test, P <0.05; Fig. 1c). However, dTRD genes are also overrepresented in SVGs in *H. vulgare, P. aphrodite*, *Z. mays*a, and some samples from *A. thaliana*. Likewise, rTRD genes are overrepresented in SVGs in *H. vulgare* and *P. aphrodite*. While some associations between duplication modes and expression profiles are data set (or species)-specific, we found consistent associations across species, indicating that duplication modes play an important role in determining a gene’s fate at the expression level.

**Fig. 1.**
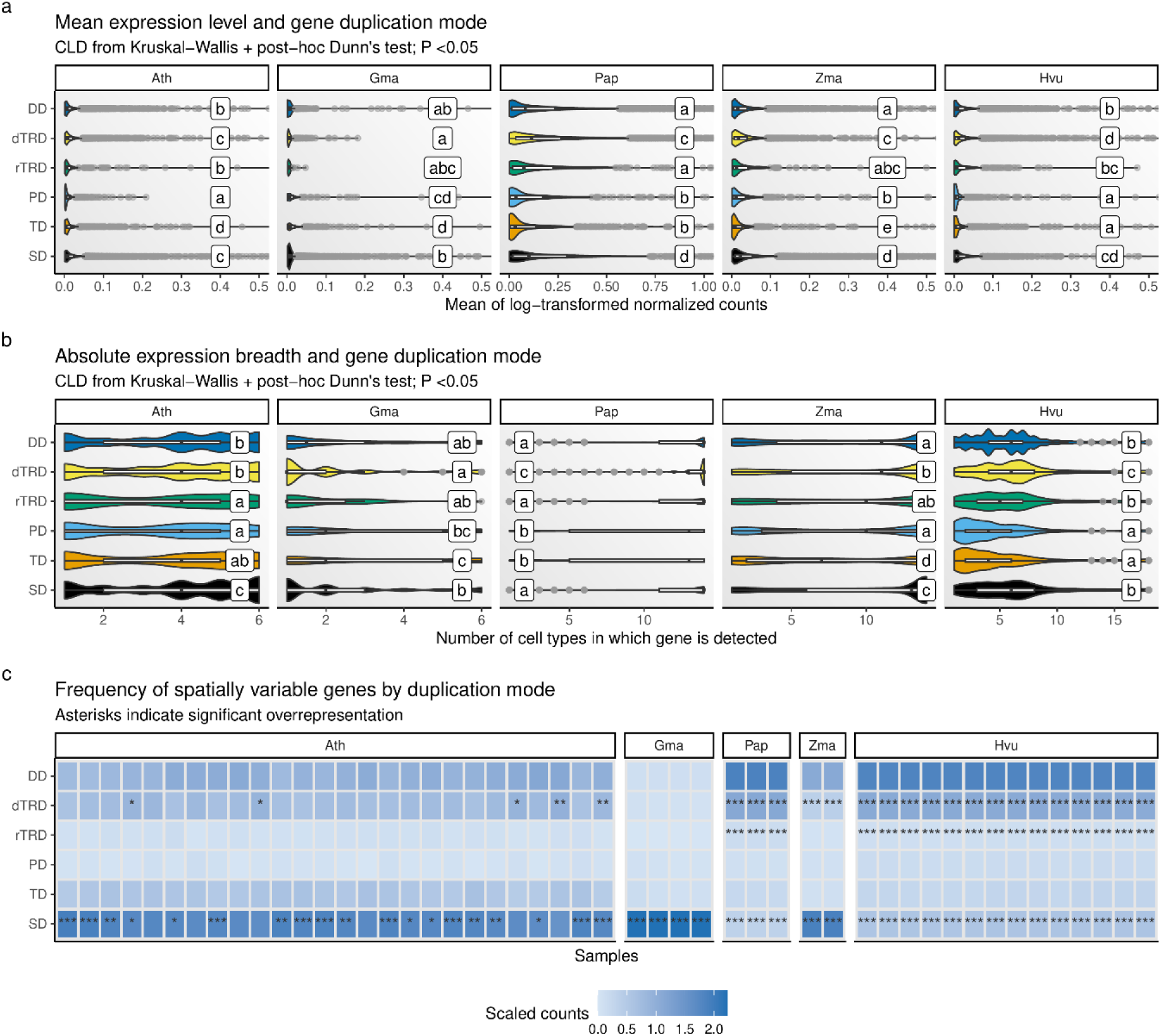
Impact of gene duplications on gene expression profiles. **a,b.** Mean expression levels and expression breadths of genes originating from different modes. Expression breadths indicate the number of different cell types in which a gene is expressed. Compact letter displays (CLDs) indicate significant differences obtained with a Kruskal-Wallis test followed by a post-hoc Dunn’s test (*P* <0.05). **c.** Frequency of spatially variable genes among genes originating from different duplication modes. Significant associations between duplication modes and spatial expression variability are indicated with asterisks, and were obtained with Fisher’s exact tests (*P* <0.05).

### Gene duplication mode impacts expression divergence in young paralogs

To investigate the impact of gene duplication modes on expression divergence between paralogs, we aggregated spots into metaspots and compared distributions of mean gene-gene Spearman’s correlations across samples (see Materials and Methods for details). We observed that expression divergence increases exponentially with time (represented by synonymous substitution rates, K_s_), as log-log regressions display a better fit compared to simple linear regressions (smaller BIC and AIC; Supplementary Fig. S2). To account for a potential confounding effect of duplication age, paralogous gene pairs were divided into age groups defined based on peaks in K_s_ distributions. Thus, SD duplicates here are likely derived from whole-genome duplications (WGDs), as they are close to K_s_ peaks. Comparisons of gene-gene correlations by duplication mode were only made for paralogs from the same age group. Gene-gene correlations were used as a measure of expression divergence, with higher correlations indicating less divergence.

We observed no differences in expression divergence for older duplicates (i.e. generated tens of million years ago, MYA) (Fig. 2a and 2b). However, younger duplicates (generated <20 MYA) display decreased expression divergence when duplicated by mechanisms that preserve *cis*-regulatory landscapes (SD, TD, and PD). For instance, younger SD and PD duplicates in soybean (originated ∼13 MYA) display less expression divergence, but no differences by duplication mode are observed for duplicates originated ∼62 MYA (Fig. 2a and 2b). Likewise, SD, TD, and PD duplicates in maize (originated ∼16 MYA) display less expression divergence, while no differences are observed for barley duplicates (originated ∼53 MYA). Collectively, our findings suggest that duplication modes impact expression divergence in an age-dependent manner. An exception exists for *A. thaliana*, whose SD duplicates display less expression divergence despite their age (∼51-72 MYA), which could be due to slower evolutionary rates, or tissue-specific differences in expression divergence rates.

**Fig. 2.**
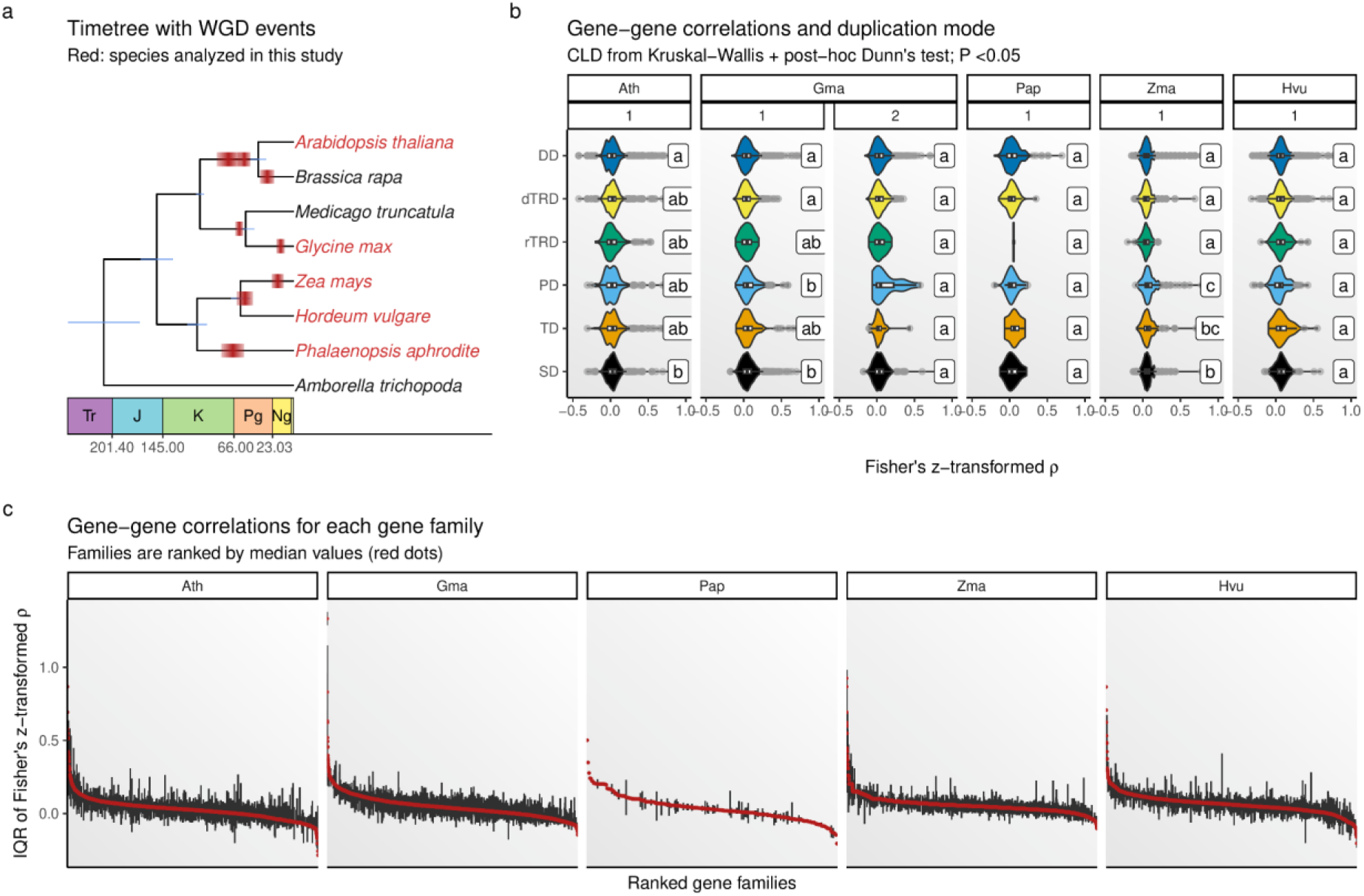
Expression divergence between paralogous gene pairs. **a.** Ultrametric species tree (a.k.a. ‘timetree’) with rectangles indicating polyploidization events. Phylogenetic positions and dates of polyploidization events were obtained from AngioWGD (Chen et al. 2024). Names of species included in this study are highlighted in red. **b.** Spearman’s correlation coefficients between paralogous gene pairs originating from different duplication modes. Numbers in facet names indicate age groups obtained from parameters of peaks in K_s_ distributions, with 1 corresponding to younger paralogs and 2 corresponding to older paralogs. Paralogs in each age group have K_s_ values near peaks in K_s_ distributions and, hence, originated by different duplication modes at around the same time of polyploidization events indicated in **a**. Compact letter displays (CLDs) indicate significant differences obtained with a Kruskal-Wallis test followed by a post-hoc Dunn’s test (*P* <0.05). **c.** Interquartile range (IQR) of correlation coefficients between paralogous gene pairs from different gene families. Red dots in the center of IQRs indicate median correlations for each gene family, which were used to arrange families in descending order. Spearman’s correlation coefficients were z-transformed.

### Expression divergence varies across gene families and functions

We hypothesized that different gene families display different rates of expression divergence, probably as a result of variability of selective constraints acting upon cis-regulatory elements of some functional groups. To test this hypothesis, we assigned paralogous pairs to gene families using delineated gene families from PLAZA (Van Bel et al. 2022), and calculated the interquartile range (IQR) of gene-gene correlations for each family. In line with our hypothesis, we observed large variations in IQRs across families, with some gene families comprising highly similar paralogs (HSPs), and some families comprising highly dissimilar paralogs (HDPs) (Fig. 2c). This finding suggests that strong selection pressures act on cis-regulatory elements of some gene families to preserve expression similarity, while more relaxed selection pressures allow genes in most other families to diverge in expression.

To better understand the functional profiles of genes in families with HSPs and HDPs, we selected the top 10% gene families with highest and lowest median gene-gene correlations, and performed an enrichment analysis for GO terms. Families with HSPs are enriched in genes associated with basic and dosage-sensitive cellular processes, including chromatin assembly, rRNA processing, regulation of transcription, splicing, and photosystem assembly. Families with HDPs are enriched in genes associated with more specialized biological processes, including flower development, auxin and abscisic acid biosynthesis, detoxification, sucrose metabolism and transport, and cell wall biogenesis and remodeling (Supplementary Table S1). Collectively, our findings highlight biological processes under stronger selection pressures, leading to increased preservation and conservation across species, and biological processes that are more likely to diversify through the subfunctionalization and neofunctionalization of genes thereof.

### Network-based expression divergence reveals potential selection pressures constraining divergence

To investigate how expression divergence between paralogs impacts sets of genes in a coordinated way, we inferred a gene coexpression network (GCN) for each species by pseudobulking and combining expression matrices (see Materials and Methods for details). In GCNs, genes with similar profiles are clustered together in so-called coexpression modules. Thus, we used co-occurrence of paralogous pairs in coexpression modules as an indication of preserved expression profiles, while paralogous pairs with genes in different modules were interpreted as having distinct (and, hence, diverged) expression profiles.

In line with our findings based on correlation distributions (Fig. 2b), we observed that most paralogous pairs display expression divergence, with relative frequencies of diverged pairs >90% in nearly all species and duplication modes (Fig. 3a). However, in most species and duplication modes, observed frequencies of diverged pairs are still smaller than the expected by chance in degree-preserving simulated networks, suggesting that selection pressures may be acting to constrain expression divergence (Fig. 3a). We found no consistent differences in network-based expression divergence by duplication mode. Module dissimilarities for diverged pairs were mostly small, indicating that, despite being in different modules (and thus displaying different expression profiles), most diverged pairs are still somewhat similar (Supplementary Fig. S3).

**Fig. 3.**
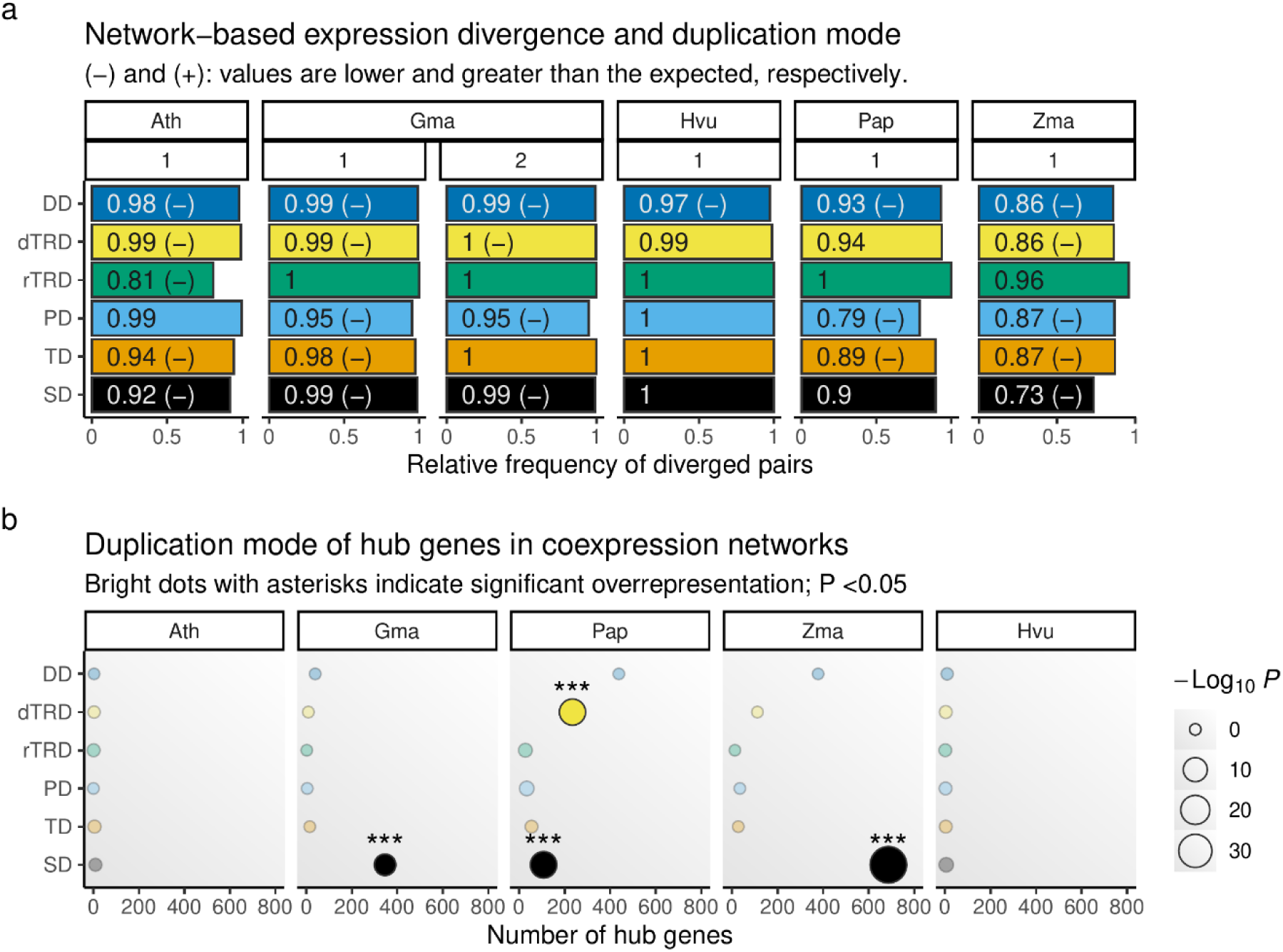
Impact of gene duplications on gene coexpression networks. **a.** Network-based expression divergence for paralogous gene pairs originating from different duplication modes. Paralog pairs are interpreted as divergent if genes therein occur in different coexpression modules. Observed frequencies of diverged pairs were compared to null distributions generated from 10,000 degree-preserving simulated networks to assess statistical significance. Numbers in facet names indicate age groups obtained from parameters of peaks in K_s_ distributions, with 1 corresponding to younger paralogs and 2 corresponding to older paralogs. **b.** Frequency of hub genes originating from each duplication mode. Asterisks indicate significant associations between hub status and duplication modes, assessed with Fisher’s exact tests (*P* <0.05).

### Gene duplication mode impacts gene connectivities in coexpression networks

We identified hubs in coexpression networks and tested for potential associations between duplication modes and gene connectivities (or node degree). We found that SD genes are enriched in hubs in *G. max*, *P. aphrodite*, and Z. mays, and dTRD genes are also enriched in hubs in *P. aphrodite* (Fig. 3b). This finding suggests that such duplication mechanisms tend to create genes with more central roles in transcriptional networks. However, the lack of association between particular duplication modes and hub status in *A. thaliana* and *H. vulgare* could indicate that such associations are species- or tissue-specific. Nevertheless, segmental duplications were significantly associated with hub status in three out of five data sets, providing stronger evidence for a potentially broader association between this duplication mode and higher connectivities.

### Pervasive asymmetric divergence among small-scale, but not segmental duplicates

To understand the evolutionary fates of paralogous pairs at the expression level, we classified paralogous gene pairs in divergence classes based on the relative expression breadths (REBs) of genes therein (see Materials and Methods for details). In short, paralog pairs are classified as displaying i. redundancy, if both genes are expressed in (nearly) the same cell types; ii. asymmetric divergence, if a gene specializes in one or a few cell types while the other remains broadly expressed; and iii. symmetric divergence, if genes are expressed in complementary, (mostly) non-overlapping cell types.

Overall, we observed that most paralogous gene pairs diverge asymmetrically, with one gene in a pair displaying high REB while the other displays low REB (Fig. 4a). This divergence pattern indicates that, in high resolution, specialization in particular cell types is the most common outcome of duplicated genes at the expression level. However, a large fraction of SD pairs display expression-level redundancy in *Z. mays* and *P. aphrodite*, as indicated by high REBs for both genes in a pair (Fig. 4a). We tested for associations between duplication modes and divergence classes, and found that paralog pairs derived from small-scale duplications (TD, PD, TRD, and DD) are overrepresented in pairs displaying asymmetric divergence in most species (Fig. 4b). Conversely, segmental duplicates are typically overrepresented in pairs displaying redundancy (Fig. 4b). Collectively, our findings indicate the duplication modes play a role in expression divergence outcomes.

**Fig. 4.**
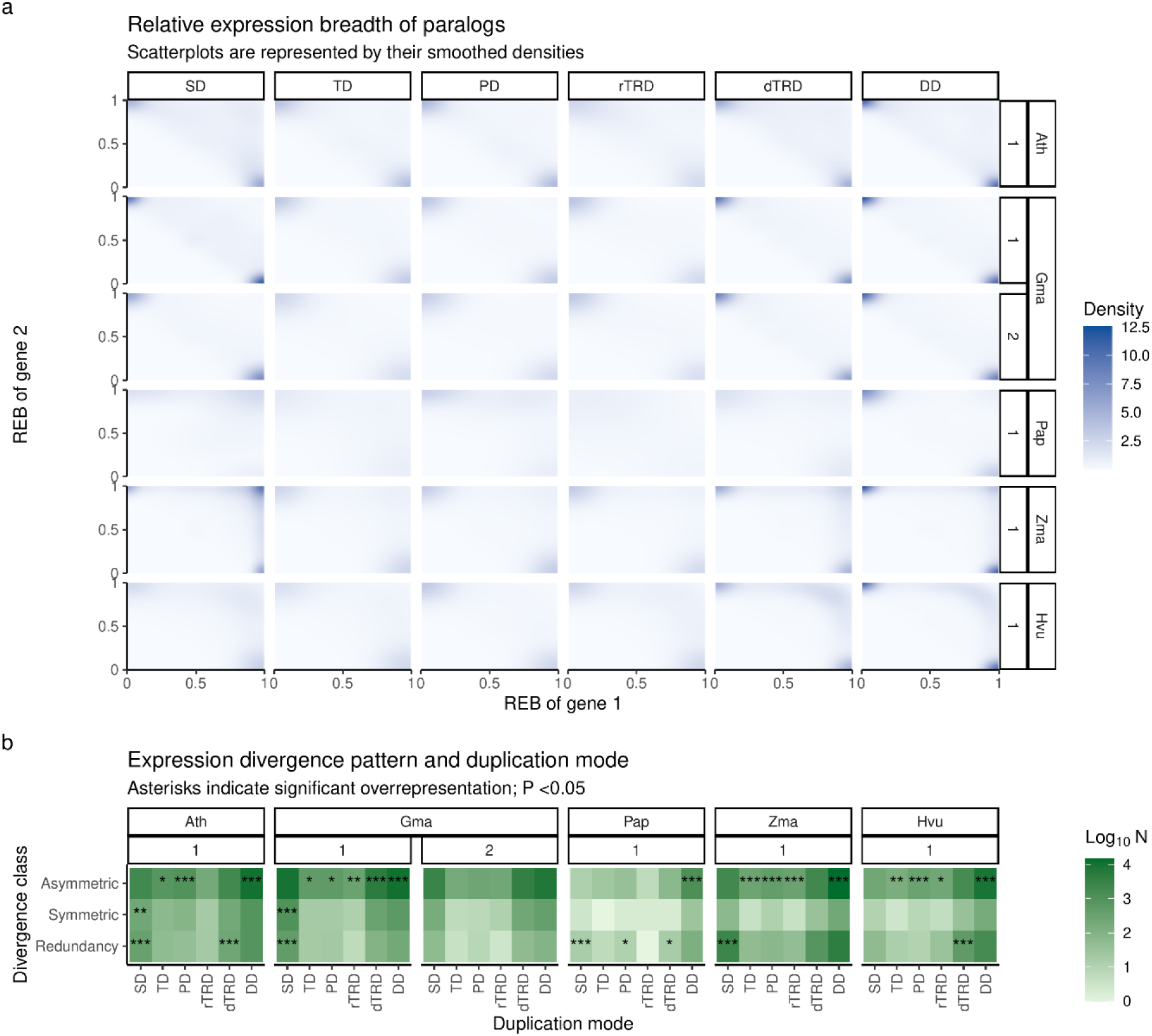
Divergence classes of paralogous gene pairs originating from different duplication modes. **a.** Relative expression breadth of genes in a paralog pair. For each paralog pair, a gene’s relative expression breadth is calculated as the ratio of the number of cell types in which it is expressed and the union of cell types in which both genes in the pair are expressed (see Materials and Methods for details). Scatterplots are represented by their smoothed densities. Numbers in facet names indicate age groups obtained from parameters of peaks in K_s_ distributions, with 1 corresponding to younger paralogs and 2 corresponding to older paralogs. **b.** Frequency of genes in each divergence class originating from each duplication mode. Asterisks indicate significant associations between divergence classes and duplication modes, assessed with Fisher’s exact tests (*P* <0.05).

## Discussion

To the best of our knowledge, this is the first study on the evolution of high-resolution expression divergence (i.e. spatially resolved, *quasi* single-cell level) following gene duplications in plants. Similarly to previous findings based on (low-resolution) bulk RNA-seq data, we demonstrate that gene duplication modes and gene functions play a major role in determining the expression-level outcome of a gene following duplication. Despite the diversity of gene duplication mechanisms, some of our findings seem to group duplication modes into two major classes based on whether or not cis-regulatory landscapes are preserved. This is highlighted by the greater expression levels and expression breadth for SD- and TD-derived genes, and the consistent association between segmental duplications and spatial variability in gene expression across species. In lotus (*Nelumbo nucifera*), greater expression levels and breadth among SD-derived genes was associated with gene essentiality, as essential genes are typically highly expressed in all plant tissues (Lloyd et al. 2015; Shi et al. 2020). However, this seems to be contradicted by the observed association between segmental duplications and spatial expression variability, because expression levels of essential genes should be stable across cell types. We hypothesize that greater and broader expression for SD- and TD-derived genes is intrinsically linked with the preservation of ancestral transcription factor binding sites (TFBS) in promoters and enhancers. Duplication mechanisms that dramatically change cis-regulatory landscapes (e.g., transposon-mediated duplications) could generate gene copies in genomic regions that have, for instance, less TFBS, or more silencing, ultimately leading to reduced and more restricted expression levels. For example, segmental duplicates in *Arabidopsis* sp. displayed less DNA methylation in promoter regions, resulting in greater expression levels (Ha et al. 2009). In *Drosophila melanogaster*, expression levels of duplicates are often greater than the expected two-fold (i.e., additive expression levels) for genes arranged in tandem, but not when the same genes are in *trans*, indicating that the tandem organization is important for non-additive increases in expression (Loehlin and Carroll 2016).

Besides the effect of shared cis-regulatory landscapes, the scale of the duplication (i.e., segmental or large-scale *versus* small-scale) plays an important role. For instance, we observed that hubs in most of the coexpression networks are enriched in SD-derived genes. Hubs are typically regarded as essential nodes for the functioning and survival of biological networks. In yeast protein-protein interaction networks, hubs were found to be enriched in essential genes (i.e., genes that lead to cell lethality when not expressed) (Jeong et al. 2001). This association was later reported in several other studies using different data sets, species, and types of biological networks, culminating in the so-called ‘lethality-centrality rule’ (Yu et al. 2004; Hahn and Kern 2005; Resendis-Antonio et al. 2005; Batada et al. 2006; Raman et al. 2014; Song et al. 2015; Almeida-Silva et al. 2020a). Essential hubs in a large-scale soybean coexpression network are involved in typically dosage-sensitive biological processes, such as nucleic acids metabolism, translation, and regulation of cell division (Tasdighian et al. 2017; Almeida-Silva et al. 2020a). As dosage-sensitive gene families are typically retained after whole-genome duplications, the association between SD-derived genes and hubs is likely due to the increased frequency of essential genes following SD (including WGD).

The scale of the duplication also clearly separates paralogs in terms of divergence classes. While duplicates originating from large-scale duplications typically undergo redundancy or subfunctionalization, small-scale duplicates often undergo neofunctionalization. Similar findings have been reported in other organisms, including *Saccharomyces cerevisiae* (Fares et al. 2013), *Arabidopsis thaliana* (Casneuf et al. 2006), *Drosophila melanogaster* (Assis and Bachtrog 2013), and grasses (Jiang and Assis 2019). In grasses, associations between small-scale duplications and neofunctionalization was hypothesized to be related to retrotransposon-derived duplications, which generate copies lacking introns and regulatory elements of their ancestral genes (Jiang and Assis 2019). However, we also observed associations between neofunctionalization and other small-scale duplication modes, such as tandem and proximal duplications, which typically preserve (although to a lesser extent) cis-regulatory landscapes. Hence, we hypothesize that differences in distal regulatory elements (e.g., enhancers) play a considerable role in the functional divergence of duplicate pairs. Alternatively, as large-scale duplications preserve genome organization, greater levels of redundancy and overlapping expression patterns among SD-derived pairs could be due to structural constraints imposed by the preservation of chromatin structure.

The effect of duplication modes and gene functions on expression divergence are mostly consistent across species, and our findings recapitulate previous findings based on bulk RNA-seq data. For instance, we demonstrate that segmental duplicates display higher correlations compared to duplicates originating from most single-gene duplication mechanisms, which is in line with previous works using large compendia of microarray and bulk RNA-seq data (Casneuf et al. 2006; Li et al. 2009; Qiao et al. 2019). Likewise, we demonstrate that duplicate pairs in dosage-dependent gene families display highly preserved expression profiles, which is likely due to stronger selection pressures to preserve stoichiometric balance, as previously suggested (Freeling and Thomas 2006; Freeling 2009; Birchler and Veitia 2012).

The consistency of findings across different species and resolutions (i.e., bulk and spatial) raises the question of whether and to what extent (duplicated) gene expression evolution is predictable. From an evolutionary lens, predicting expression divergence following duplications can help understand how the rewiring of gene networks shaped the origin and diversification of traits, similarly to previous works on flowering (Calderwood et al. 2021), nodulation (Libourel et al. 2023), organogenesis (Julca et al. 2021), and response to changing environments (Hamann et al. 2021). From a biotechnological perspective, predicting gene expression evolution could help overcome barriers in crop breeding resulting from paralog diversification. For instance, gene duplications and subsequent paralog diversification within key domestication-related gene families in *Solanum* sp. hinders genotype-to-phenotype predictability, limiting knowledge transfer between related crops (Benoit et al. 2025).

While gene expression evolution is greatly shaped by a gene’s duplication mode and functions, our findings highlight that time is an important (and often overlooked) variable. We demonstrate that paralog pairs originating from different duplication modes display differences in expression divergence, but such differences only exist for young duplicates, indicating that the effect of duplication mode is age-dependent. This is in line with our previous finding that WGDs have a significant impact on plant gene regulatory networks (GRNs), increasing the frequency of genetic circuits called GRN motifs, but WGD-derived motifs are lost over time (Almeida-Silva and Van de Peer 2023). Likewise, WGD-derived duplicates in soybean display greater levels of expression preservation, but only for those arising from the most recent WGD at ∼13 million years ago (Almeida-Silva et al. 2020b).

By using high-resolution, spatially resolved transcriptomic data, this study advances our understanding of the tempo and mode of gene expression evolution following gene and genome duplications. Since our findings here are similar to previous findings based on bulk RNA-seq data, a question that remains open is whether patterns of expression divergence are universal and independent in both low and high resolution (i.e., across tissues *versus* across cell types). Specifically, it remains unclear whether expression divergence follows an order in terms of resolution, with duplicates first diverging across cell types in a tissue and then across multiple tissues in an entire plant (or vice versa), or if duplicates diverge across cell types and across tissues independently. Addressing this question requires comparing bulk and spatial transcriptomic data from the same species under the same experimental conditions, which we believe will be possible in the near future as new data become available and technologies advance.

Despite the high resolution and throughput of spatial transcriptomics technologies, tissue capture areas are small, with slides of 6.5 mm x 6.5 mm for the 10x Visium, for instance. This limitation might raise questions as to whether our observations are replicable in other tissues and conditions. Since our findings are consistent across independent data sets, we believe they broadly represent the biology of gene expression divergence within tissue compartments. While only a few spatially resolved transcriptomes exist for plants, we expect a steep increase in the number of spatial transcriptomics studies in the coming years. As more data become available, an expanded version of this study will be valuable to assess whether our findings here are broadly valid, or if exceptions exist for particular tissues or species.

### Conclusions

In this paper, we analyzed five spatially resolved plant transcriptomes to investigate the evolution of gene expression following gene and genome duplications. We found that genes originating from different duplication modes display differences in expression levels, expression breadth, spatial variability, and number of coexpression partners. Duplication mechanisms that preserve cis-regulatory landscapes tend to lead to more preserved expression profiles, but differences by duplication mode disappear over time. We also observed that genes originating from large-scale (or segmental) and small-scale duplications undergo different functional divergence outcomes. Our findings align with previous studies based on low-resolution (bulk) expression data, suggesting that patterns of expression divergence following gene duplications are universal within the same tissue and across different tissues.

## Materials and Methods

### Obtaining and preprocessing spatial transcriptomic data

We manually curated the literature to find spatially resolved plant transcriptomes obtained with high-throughput, sequencing-based technologies. Data sets representing Arabidopsis leaves (Xia et al. 2022), *Phalaenopsis* Big Chili inflorescences (Liu et al. 2022), and maize ear primordia (Wang et al. 2024) were obtained from STOmicsDB (Xu et al. 2024), while data sets representing barley seeds (Peirats-Llobet et al. 2023) and soybean nodules (Liu et al. 2023) were obtained from original publications. Data from H5AD files were imported with the R package zellkonverter (Luke Zappia, Aaron Lun), and stored as *SpatialExperiment* objects (Righelli et al. 2022). Data were filtered to remove genes with zero counts across all spots, spots with zero counts, and genes with less than 1 count in ≥99.9% of the spots. Filtered data were preprocessed to correct for library size differences and compute log-transformed normalized counts with the R package scater (McCarthy et al. 2017).

### Obtaining classified duplicated gene pairs and genes

Duplicated genes and gene pairs classified by duplication mode were obtained from doubletroubledb (Almeida-Silva and Van de Peer 2025). As *Phalaenopsis aphrodite* was not included in this database, we identified and classified its duplicated genes and gene pairs with doubletrouble (Almeida-Silva and Van de Peer 2025) using *Amborella trichopoda* as an outgroup. Duplicated gene pairs were classified as originating from segmental duplications (SD, duplication of large chromosomal regions, whole chromosome(s), or the whole genome), tandem duplications (TD, duplicated copy adjacent to the ancestral gene), proximal duplications (PD, duplicated copy near the ancestral gene, but separated by a few genes), retrotransposed (rTRD, insertion of retrotransposed gene), DNA transposed (dTRD, duplicated copy generated by the activity of DNA transposons), and dispersed duplications (DD, none of the above). Functional annotation and gene family assignments were obtained from PLAZA 5.0 (Van Bel et al. 2022).

### Calculation of substitution rates and age-based grouping of paralogous gene pairs

Synonymous and nonsynonymous substitution rates (K_s_ and K_a_, respectively) were calculated with the γ-MYN model (Wang et al. 2009) implemented in the R package doubletrouble (Almeida-Silva and Van de Peer 2025). Coding sequences (CDS) for the longest isoform of each gene were obtained from Ensembl Plants release 59 (Yates et al. 2022) and Orchidstra 2.0 (Chao et al. 2017). After removing pairs with K_s_ >2 to avoid saturation at greater K_s_ values, we used doubletrouble (Almeida-Silva and Van de Peer 2025) to infer peaks in K_s_ distributions with Gaussian Mixture Models (GMMs), with pre-defined number of components based on literature evidence (Chen et al. 2024). Parameters of Gaussian components (i.e., K_s_ peaks) were used to group paralogous gene pairs into age groups, defined based on the mean ±2 standard deviations of each component.

### Inference of spatially variable genes and overrepresentation analyses

Spatially variable genes were inferred with DESpace (Cai et al. 2024) using annotated cell types as spatial clusters. Curated spatial domain annotations were obtained from original publications. Genes were considered spatially variable if FDR <0.05. Overrepresentation analyses were performed with the R package HybridExpress (Almeida-Silva et al. 2024).

### Measuring expression divergence between paralogous gene pairs

We measured expression divergence between paralogous pairs using three metrics implemented in the R package exdiva (Almeida-Silva and Van de Peer 2024), namely i. Spearman’s correlations coefficients between gene pairs, ii. co-occurrence in coexpression modules, and iii. dissimilarities between module eigengenes. To avoid zero-inflated correlation distributions due to the inherent sparsity of spatial transcriptomics data, we aggregated counts for spots into metaspots, as previously suggested (Morabito et al. 2023). To select a proper number of spots in each metaspot considering the dimensions of plant cells, we simulated increasingly large metaspots (*N* = 10, 20, 30, 40, and 50 spots), and inspected distributions of gene-gene correlations (Supplementary Fig. S2). Based on a trade-off between the reduction of noise arising from sparsity and preservation of expression resolution, we used metaspots of size 30. Following previous papers (Gu et al. 2002; Makova and Li 2003; Ganko et al. 2007), Spearman’s correlation coefficients were transformed using Fisher’s z-transformation, defined as:

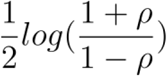

### Inference and analysis of gene coexpression networks

For each sample, we performed pseudobulk aggregation of counts (‘sum’ method) by spatial domain using the R package scuttle (McCarthy et al. 2017). Pseudobulk count matrices from the same species were combined, resulting in a single multi-sample pseudobulk matrix per species. Pseudobulk matrices were used to infer a gene coexpression network for each species using the R package BioNERO (Almeida-Silva and Venancio 2022). Signed hybrid networks (i.e., negative correlations set to zero) were inferred using Spearman’s correlation coefficients as a similarity metric. The R package exdiva (Almeida-Silva and Van de Peer 2024) was used to classify paralogous pairs based co-occurrence in coexpression modules. Paralogous pairs were classified as ‘preserved’ if both genes were in the same module, and ‘diverged’ otherwise. To assess the significance of the frequencies of diverged pairs, we generated 10,000 degree-preserving simulated networks through node label permutation, and compared observed frequencies to null distributions. Dissimilarities between module eigengenes (i.e., first principal component of the expression matrix for a module, summarizing its expression profile) were used as module distance measures. Hub genes were identified with the function *get_hubs_gcn()* from the R package BioNERO (Almeida-Silva and Venancio 2022).

### Classification of paralogous gene pairs into divergence classes

Paralogous gene pairs were classified into divergence classes based on the relative expression breadths (REBs) of their genes, similarly to a previous work (Casneuf et al. 2006). For each paralogous gene pair P with genes A and B, the relative expression breadth for gene A is defined as:

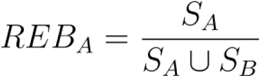

where S_A_ represents the set of cell types in which gene A is expressed, and S_B_ represents the set of cell types in which gene B is expressed. Paralogous gene pairs were classified as displaying i. redundancy (i.e., both genes with REB >0.7), ii. asymmetric divergence (i.e., one gene with REB >0.7, and the other gene with REB <0.3), or iii. symmetric divergence (i.e., both genes with REB between 0.3 and 0.7).

## Acknowledgements

YVdP acknowledges funding from the European Research Council (ERC) under the European Union’s Horizon 2020 research and innovation program (No. 833522). YVdP and FA-S acknowledge funding from Ghent University (Methusalem funding, BOF.MET.2021.0005.01).

## Author contributions

FA-S and YVdP conceived the study; FA-S collected and analyzed the data; FA-S and YVdP wrote the manuscript. All authors read and approved the manuscript.

## Data availability statement

To ensure full reproducibility, all code and data used in this manuscript are available in a GitHub repository at https://github.com/almeidasilvaf/PlantSpatialDiv, and archived at Zenodo (Almeida-Silva 2025).

---

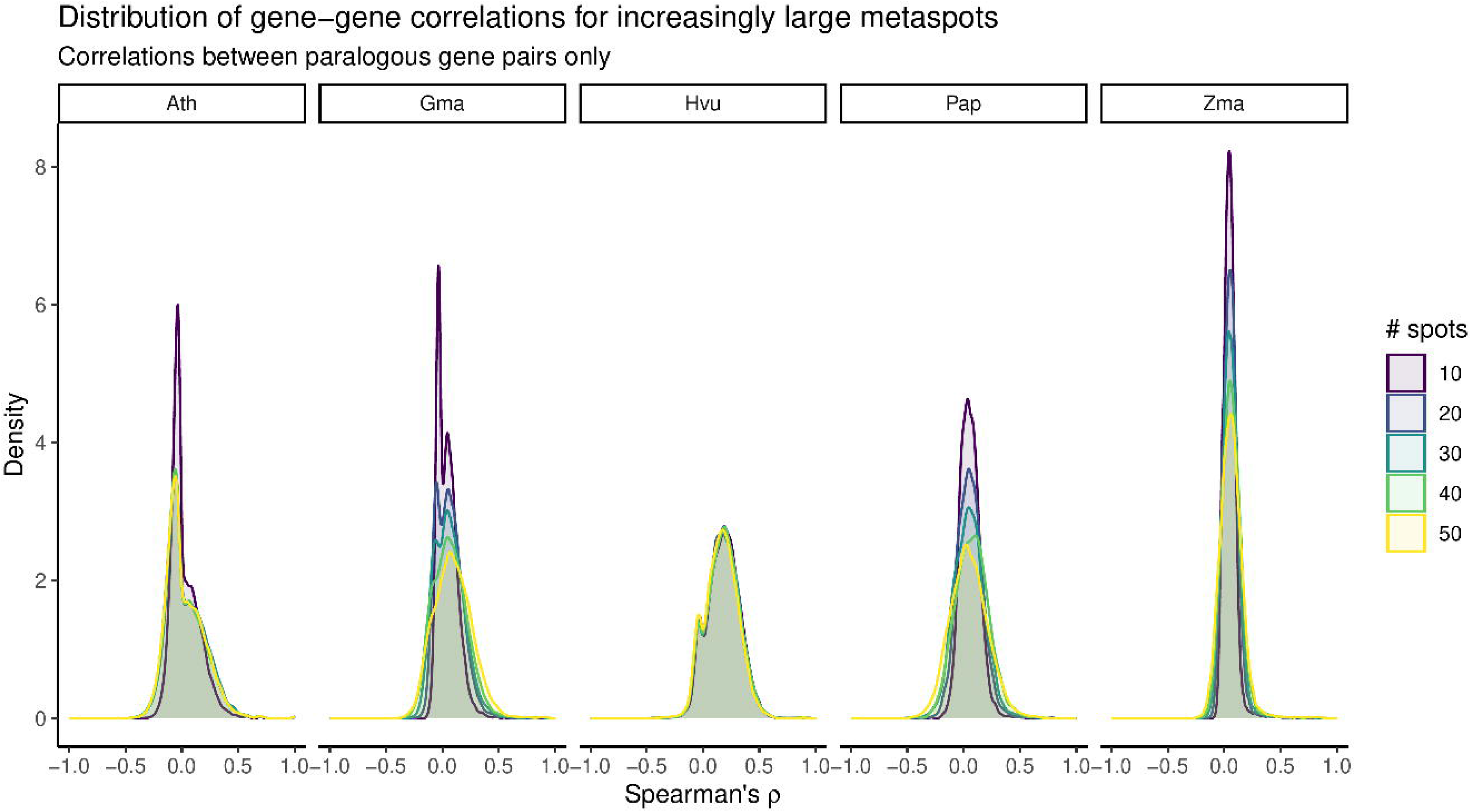

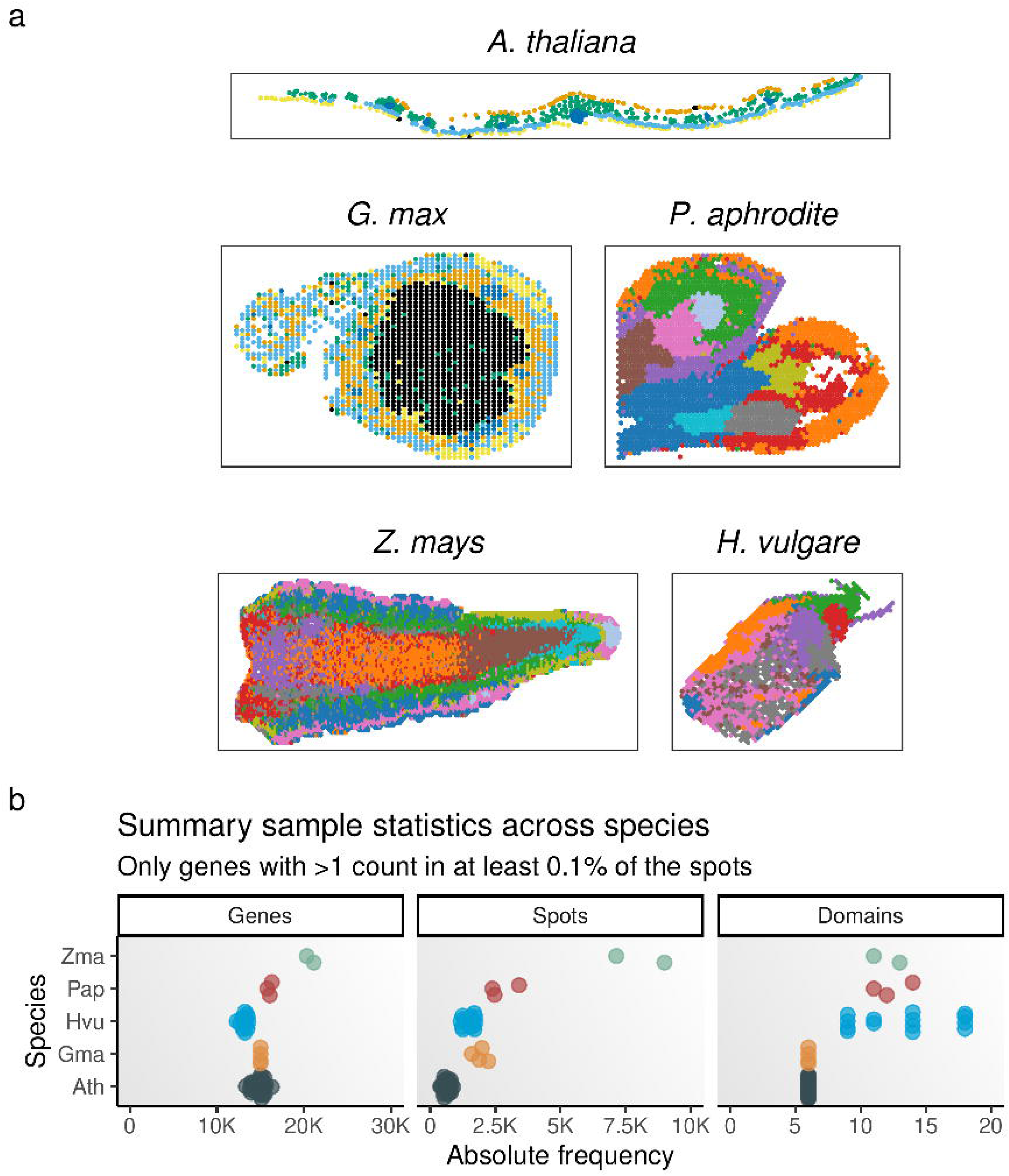

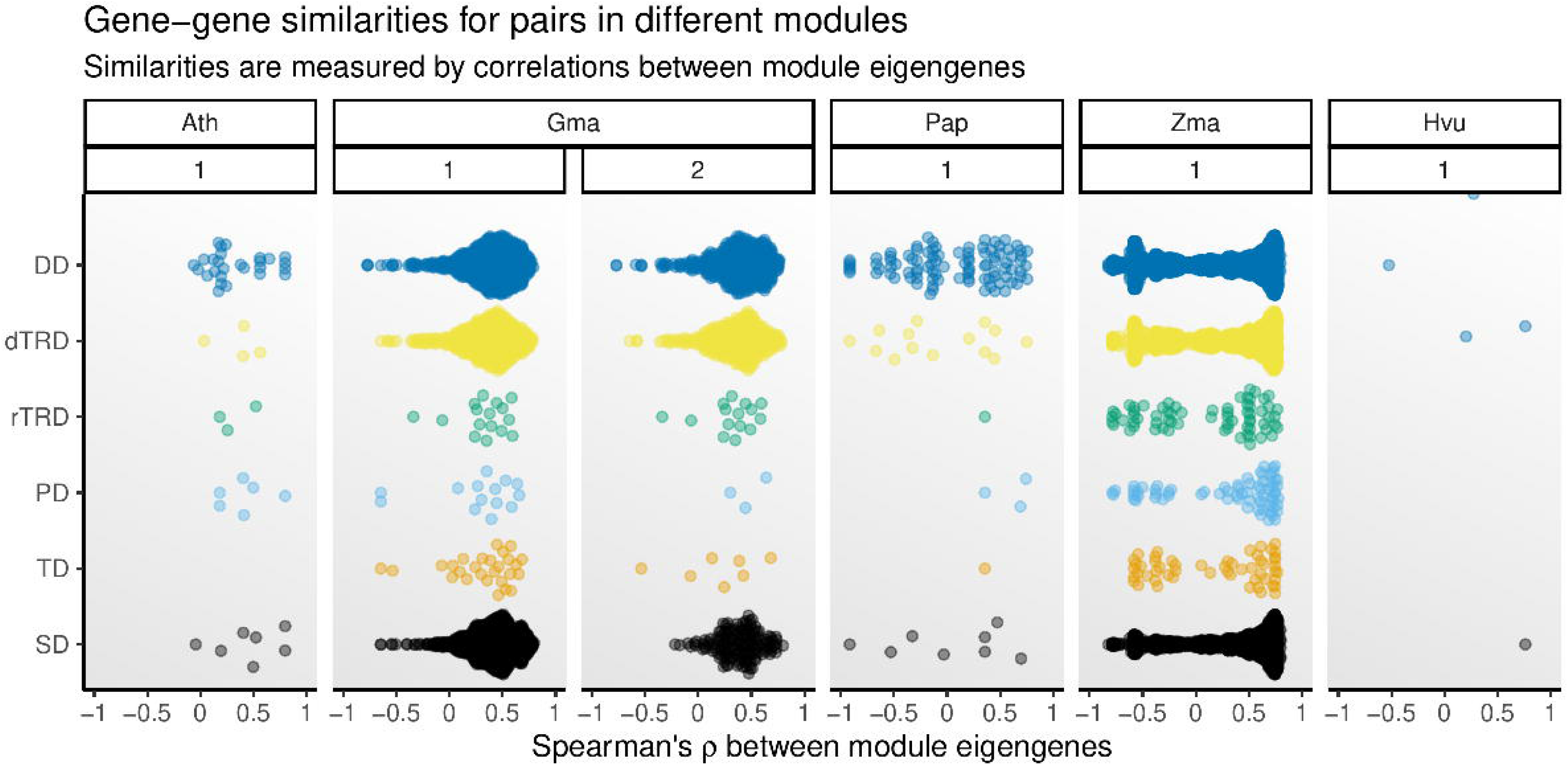

