## Supplementary Figures for "Gene expression divergence following gene and genome duplications in spatially resolved plant transcriptomes"

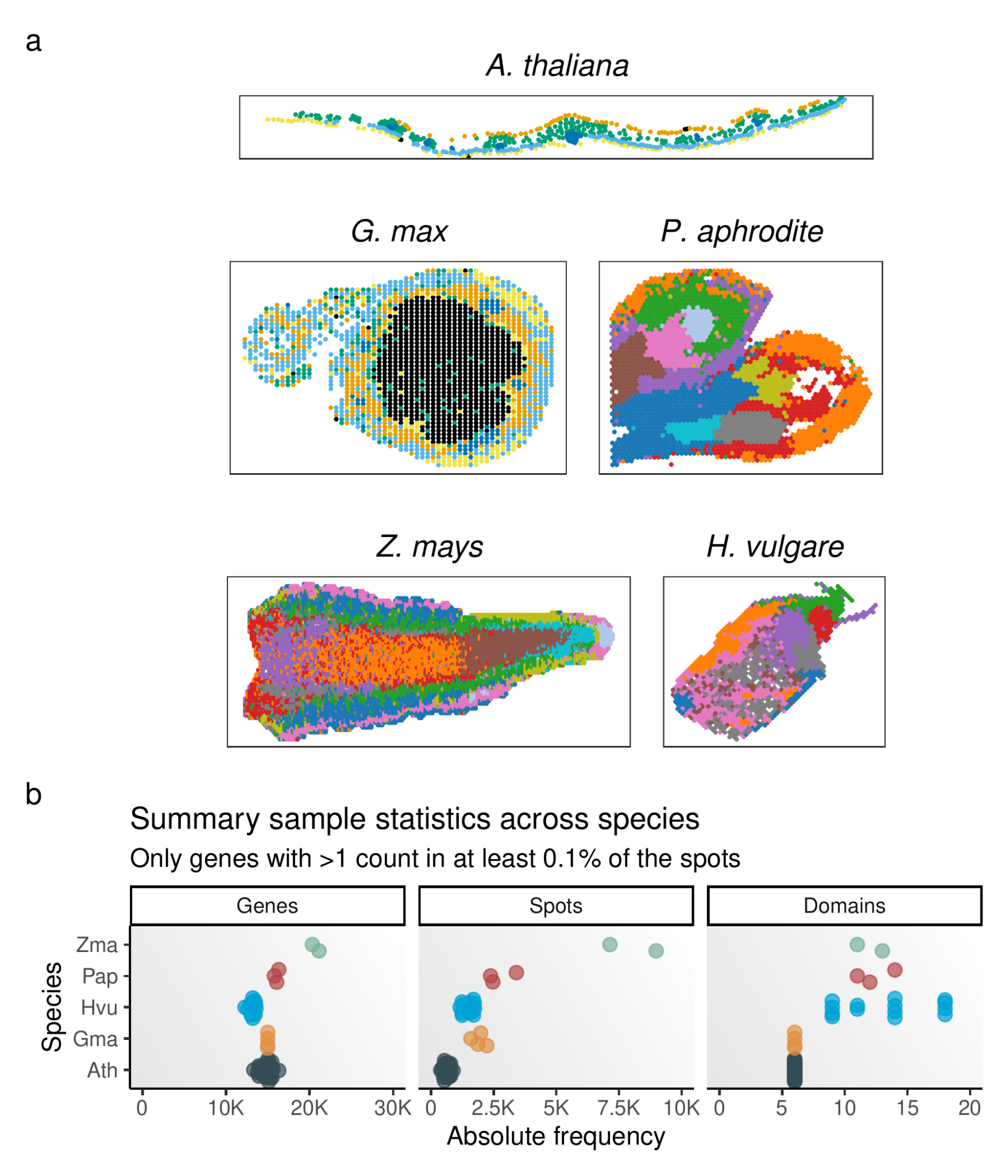

**Fig. 1. Data sets used in this study.** **a.** Representative slides for each data set. Data sets represent Arabidopsis leaves, soybean (*Glycine max*) nodules infected with rhizobia, inflorescences of Phalaenopsis Big Chili, developing maize (*Zea mays*) ears, and germinating barley (*Hordeum vulgare*) seeds. **b.** Summary statistics for each sample. Statistics shown are the number of genes, number of spots, and number of spatial domains (obtained from original publications).

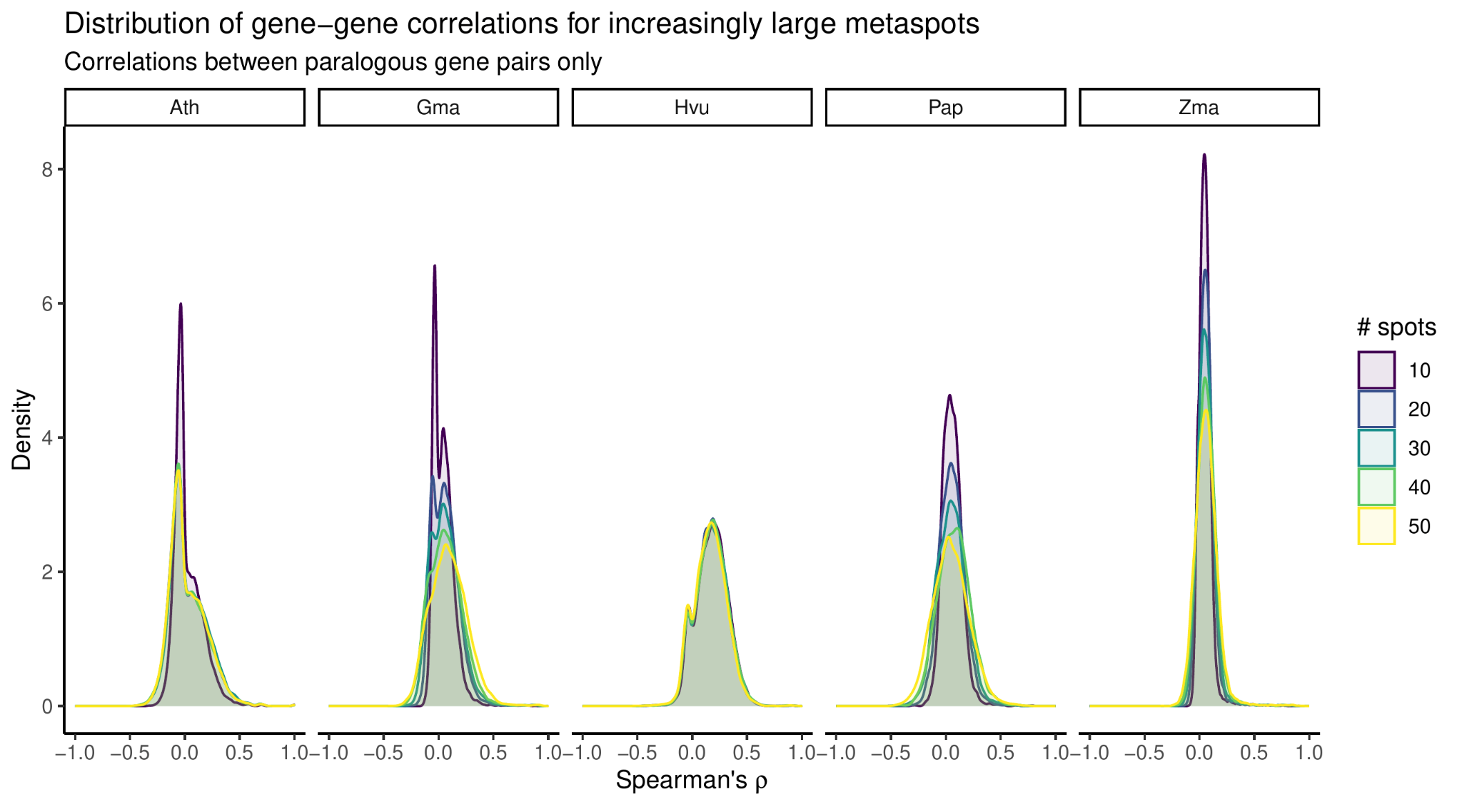

**Fig. 2. Simulation of gene-gene correlation distributions for increasingly large metaspots.** For each sample, spots were combined into metaspots using k-means clustering on a matrix of spatial (x and y) coordinates. For increasingly large metaspots, density plots indicate how metaspot aggregation reduces zero inflation arising from sparsity in correlation distributions.

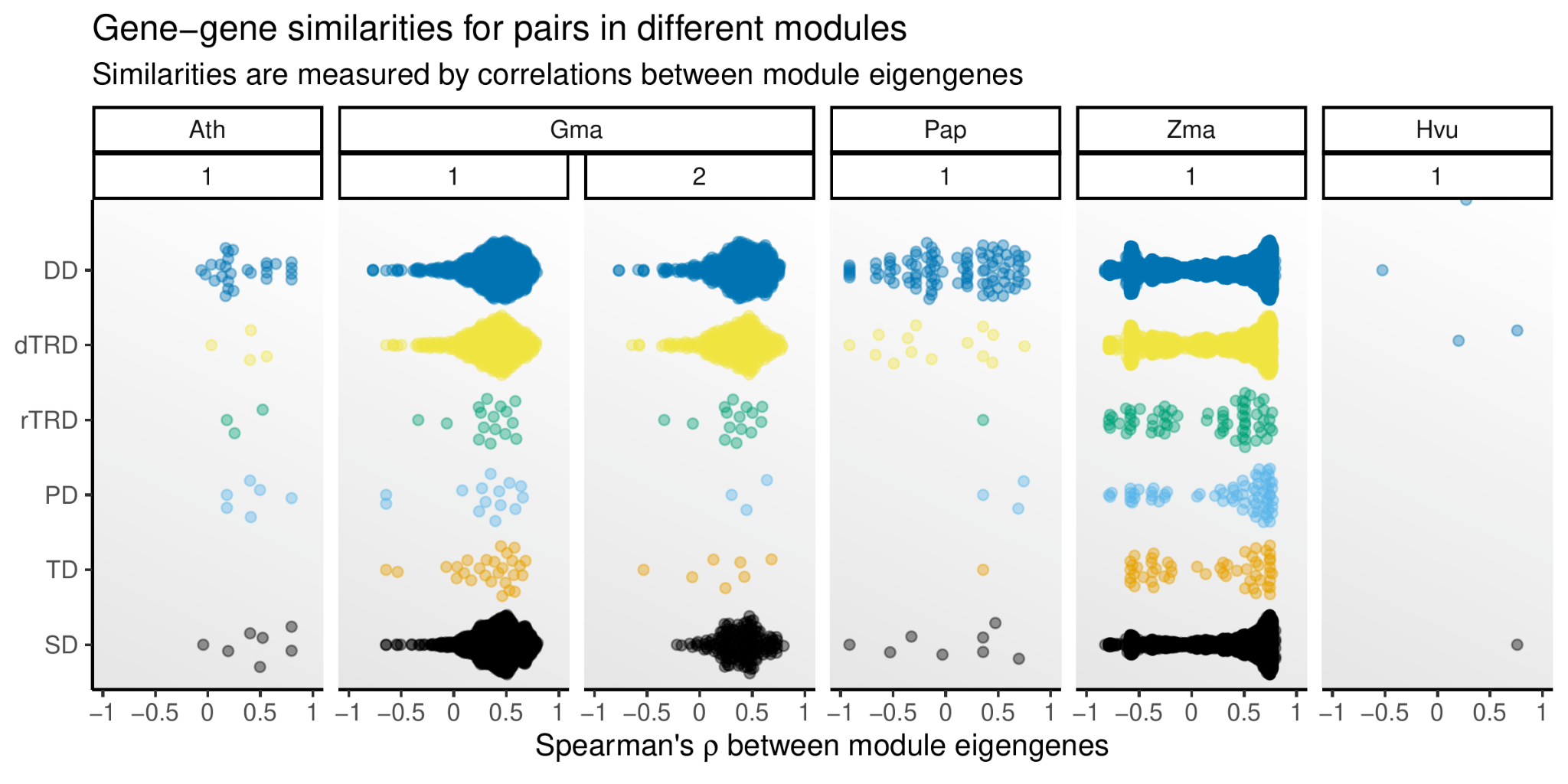

**Fig. 3. Correlation between module eigengenes for duplicate pairs in different coexpression modules.** Module eigengenes are the first principal component of the expression matrix for a module, representing a summary of the module’s expression profile.
